# Understanding the emergence of the influenza A/H3N2 K subclade in its historical and evolutionary context

**DOI:** 10.64898/2026.05.21.726823

**Authors:** Kieran Dee, Ryan M Imrie, Oscar A MacLean, Laura Mojsiejczuk, Ewan W Smith, Savitha Raveendran, Kieran Lamb, Hanting Chen, Verena Schultz, Ziping Wang, Sarah K Walsh, Junsen Zhang, Edward K Hutchinson, Brian J Willett, Emma C Thomson, Joseph Hughes, David L Robertson, Christopher J R Illingworth, Pablo R Murcia

**Affiliations:** MRC-University of Glasgow Centre for Virus Research, University of Glasgow, Glasgow, United Kingdom

## Abstract

The emergence in 2025/26 of the influenza A/H3N2 K substrain (H3N2/K) was the cause of significant public health concern. This genetically divergent virus was assessed to have a strongly decreased reactivity to contemporary vaccine strains. Respectively prolonged and early influenza seasons in the Southern and Northern Hemispheres contributed to concerns about vaccine efficacy. Here we retrospectively assessed the genetic and antigenic properties of this virus, combining epidemiological surveillance data, computational antigenic analysis, and serological data using samples from a well-stratified UK cohort. In contrast to initial indications, we found that despite the genetic distinctiveness of H3N2/K the virus had undergone limited antigenic change, suggesting that its emergence was instead the result of selection for non-antigenic properties. We confirmed previous results showing that contemporary vaccines produced an enhanced neutralising response to H3N2/K but, in a stratified serological analysis, showed that responses to the J and K substrains were age-dependent, largely driven by patterns of vaccination. Our results have implications for antigenic surveillance and for public communication strategies in future influenza seasons.

## Introduction

The spread of human influenza strains is shaped by epidemiological responses, virus evolution, and complex immunodynamics. Escape from strain-specific immunity is a key driver of virus evolution^1,2^, with cross-reactive antibodies and T-cell responses also exerting selective pressure^3,4^. Virus evolution occurs within a set of constraints, including the need to maintain protein functions such as specific receptor binding^5^, and protein structural constraints, with for example protein stability^6,7^ and epistatic effects^8,9^ influencing evolutionary trajectories. These factors create a pattern of regular selective sweeps driven by changing antigenically dominant strains from year to year^10^, whereby circulating viruses descend from a relatively recent common ancestor.

In 2025 the rapid emergence of a divergent influenza A/H3N2 strain was noted first in the USA^11^. This was subsequently named subclade K (H3N2/K) in 2026 following its global spread^12^. When compared to its closest ancestors (members of subclade J), H3N2/K included several nonsynonymous mutations in the hemagglutinin (HA) segment, some of which occurred at previously characterised antigenic sites^13^. Of note, the emergence of H3N2/K was associated with a prolonged influenza season in Australia and New Zealand^12^ and an early start of the influenza season in the UK^14,15^.

Serology studies using ferret antisera on hemagglutination inhibition (HI) assays suggested that_H3N2/K was antigenically distinct from related strains, with ferrets exhibiting a 32-fold reduction in HI activity towards the strain relative to that towards vaccine strains used in the 2024/25 and 2025/6 winter seasons (A/Thailand/08/2022 and A/Croatia/10136RV/2023 respectively) ^16^. The early onset of the 2025/26 influenza season in the UK contributed to concerns about the emergence of a highly mutated virus with reduced population immunity. Media reports unhelpfully popularised the description of the strain as a ‘superflu’^17,18^. However, while the K subclade became globally predominant^19^, an epidemiological analysis of UK case numbers indicated that a scenario involving substantial immune escape was unlikely^20^. Importantly, studies of vaccine efficacy suggested that, despite the large reported antigenic novelty of the virus, existing vaccines were effective in individuals under the age of 65^21,22^.

The human immune response to influenza infection at a population scale is a critical determinant of the spread and severity of a seasonal epidemic. Serological studies using human antisera from US and UK populations suggested that H3N2/K was antigenically distinct from previously circulating viruses, but with a less dramatic difference between strains than suggested by the original ferret data^23–25^. These studies also confirmed the effectiveness of vaccination in increasing immune protection against H3N2/K. A more recent analysis used sera from the US and Asia to evaluate a library of H1 and H3 haemagglutinin proteins, again highlighting partial, rather than dramatic, immune escape in the H3N2 K subclade[32].

Noting the mismatch between some early public health messaging and the actual impact of H3N2/K, we here assess, in retrospect, the emergence of this dominant subclade. Combining surveillance, computational and experimental serology methods, we investigate the mismatch between the apparent genetic and antigenic novelty of the K subclade and its actual epidemiological impact. Computational modelling shows that, in the context of recent genetic and antigenic virus evolution, H3N2/K was a significant outlier, exhibiting a low amount of antigenic change with respect to its genetic novelty. Data from human sera showed a considerably smaller difference between representative J and K subclade viruses than did the initial assessment of ferret sera. We consider the value of different approaches in the early assessment of evolved seasonal influenza viruses.

## Results

### Surveillance data

Community surveillance data collected systematically from primary care facilities in Scotland illustrated the epidemiological impact of H3N2/K emergence. In common with data collected elsewhere in the UK^26^, data local to Scotland showed the 2025/6 season to have an earlier peak in the incidence of influenza A and in the fraction of test rates that were positive compared to other recent seasons, albeit that the total number of reported cases were not noticeably higher than in previous seasons (Figure 1). In this sense, 2025-6 was not a particularly atypical year for influenza A. Concerns of a particularly severe influenza season in the 2025/6 Northern Hemisphere winter season were not realised^17,18^.

**Figure 1:**
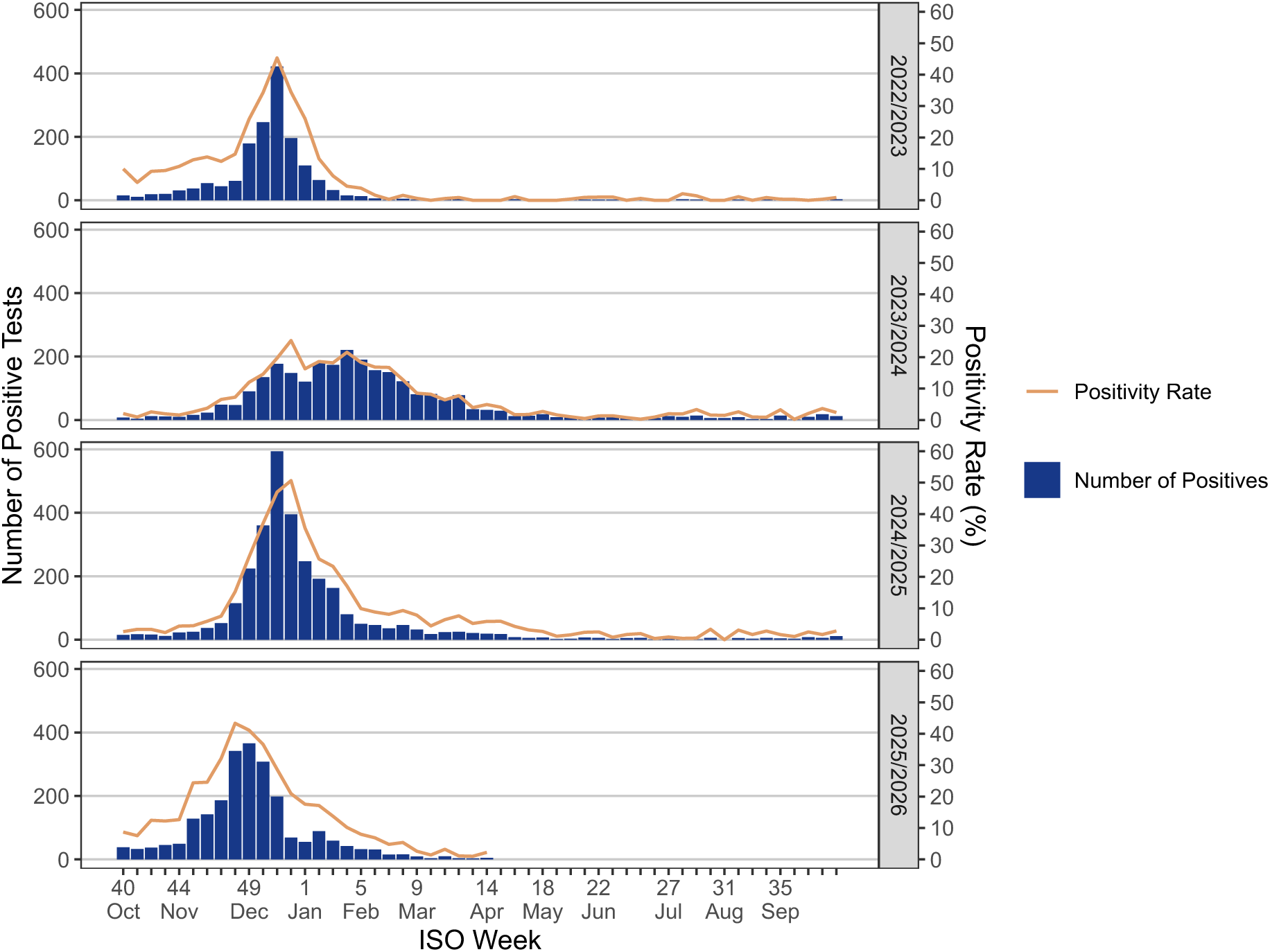
Positive tests and positive test rates in Scotland for cases of influenza. **A.** Blue bars show numbers of positive tests for influenza A each week, while the orange line shows the proportion of tests conducted that were positive for influenza A. Graphs were generated using publicly available information reported by Public Health Scotland.

### Viral genomic analysis

A phylogenetic analysis of influenza A/H3N2 HA protein sequences available from 1997 to 2026 (rooted with earlier sequences, Figure S1) showed the HA segment of H3N2/K to contain a greater than average number of non-synonymous substitutions relative to viruses from the 2024/25 season (Figure 2A), although not enough for the strain to be a significant outlier with respect to a distribution fitted to the previous data (p=0.07). The phylogeny identified multiple historical non-synonymous selective sweeps in the primary lineage of the viral population (Figure S1). The eight HA substitutions observed between consensus sequences in 2025 and 2026 was the second highest number of per-year changes since 2013, the exception being the 2020/1 season, where ‘lockdowns’ and associated measures taken in response to the COVID-19 pandemic produced uncharacteristic viral dynamics in influenza populations[35]. A negative binomial distribution describing numbers of HA substitutions per year was fitted to data from seasons up to 2024/5 (Table S1). Relative to this, the number of substitutions observed in 2025/6, reflecting the emergence of H3N2/K, was not significantly high at the 5% level. The 2020/1 season was the sole outlier in this analysis (p=0.019).

**Figure 2:**
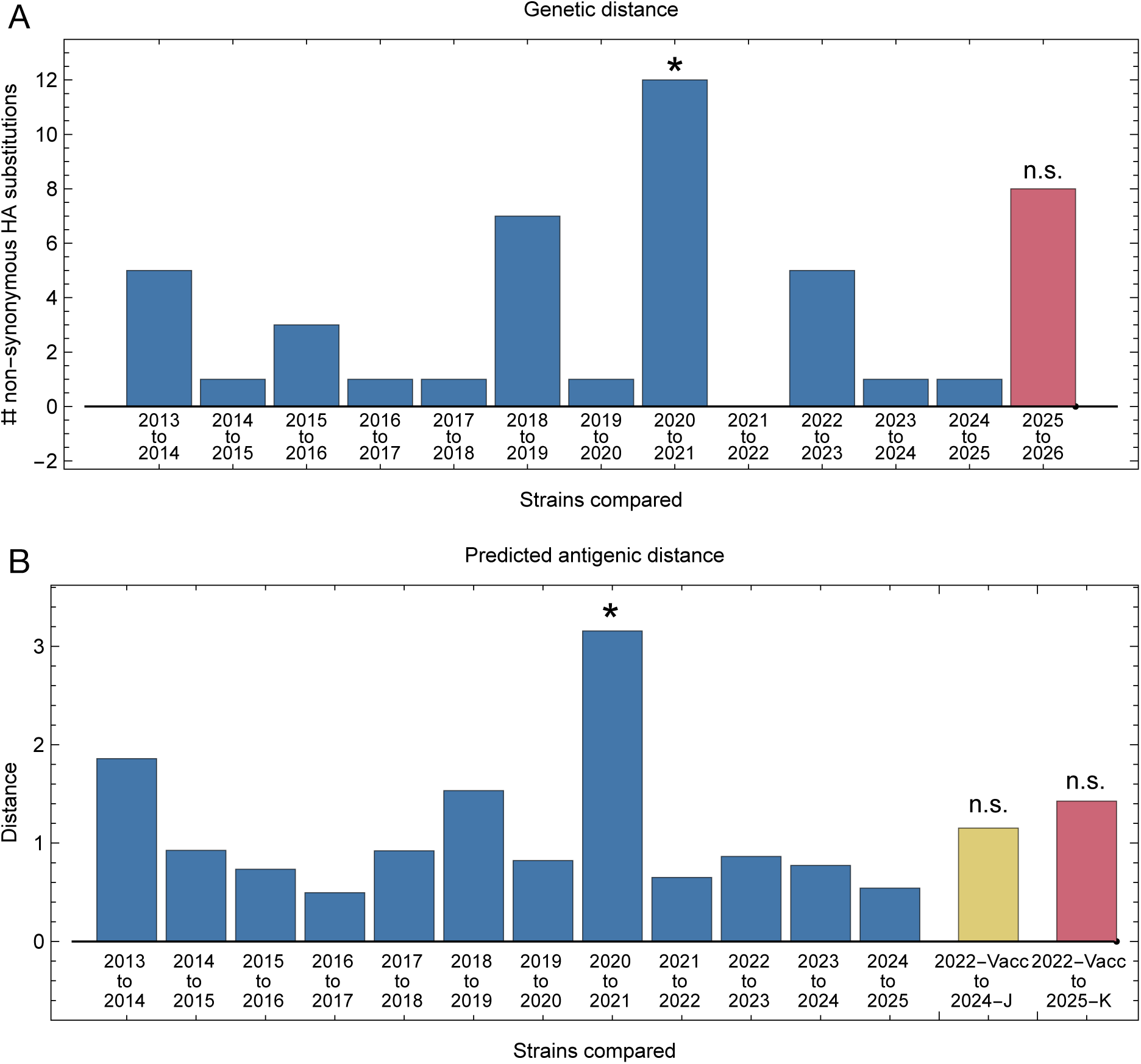
Computational analysis of the influenza A/H3N2 K subclade. **A.** Timings of fixation events observed on a phylogenetic tree describing relationships between HA viral segments presented as a count of events per year. The switch to the K subclade occurred between 2025 and 2026. Significance was calculated on the basis of a negative binomial model trained on the data from 2013/14 to 2024/25 (* p<0.05) **B.** Forecast antigenic distances generated by the PLANT evolutionary model. Distances in blue show distances in projected antigenic space between the mean locations of sequences in subsequent years. The yellow and red bars show predicted distances between the 2024-5 vaccine strain and representatives of the J and K subclades. Significance was calculated using a negative binomial model fitted to the data from 2013/14 to 2024/25. Neither distance is statistically significant in the context of previous events.

### Antigenic properties of the H3N2/K strain: Forecasting

We introduce notation to denote three viruses that were important to our evaluation. We considered the 2024/25 vaccine strain A/Thailand/08/2022, here denoted 2022-Vacc, alongside representatives of the J and K subclades of influenza A/H3N2, namely A/England/5160964/2024, denoted 2024-J, and A/England/01837755/2025, denoted 2025-K.

A forecast of antigenic differences based upon antigenic cartography integrated with a protein language model^27^ suggested that H3N2/K was antigenically distinct from the 2024/25 vaccine strain, but in a manner consistent with previous year-to-year differences between influenza A/H3N2 strains (Figure 2B). The language model maps HA1 protein sequences into a three-dimensional antigenic space (Figure S2). Mean distances in this space between strains from previous years were calculated, alongside strain-specific distances between 2022-Vacc, 2024-J, and 2025-K. Year-to-year differences in antigenic phenotype were used to fit a lognormal distribution of annual antigenic change (Table S2). The strain-specific distances from 2022-Vacc to 2024-J and 2025-K were not significantly large with respect to this distribution, with 2025-K, at 1.42 units from 2022-Vacc, not forecast to provide substantially more immune escape than did 2024-J (1.15 units). These distances correspond to roughly 2.7 and 2.2-fold decreases in immune response with respect to the vaccine strain.

Under this metric, the calculated difference in escape between the J and K representative strains, of around 0.3 units, was small. Similar results were obtained when we replaced 2022-Vacc with the 2025/26 winter vaccine strain A/Croatia/10136RV/2023 provided similar results (Figure S3). The latter vaccine strain was forecast to be slightly closer in antigenic space to the 2024-J and 2025-K viruses (0.60 and 0.97 units respectively). Again, 2025-K was forecast to provide only marginally more immune escape than 2024-J. We note that, in common with the genetic data, the forecast antigenic change for the 2020-1 season was unusually high (p=0.012).

Further analysis identified H3N2/K as an outlier with respect to its combined genetic and forecast antigenic properties (Figure 3). Fitting a linear model to data between 2013 and 2025 showed a clear positive relationship between the number of nonsynonymous substitutions in HA and the forecast antigenic distance (R^2^=0.87). Evaluating the 2025/26 transition in the context of this model suggested it to have a significantly low forecasted antigenic distance given the number of observed nonsynonymous substitutions (p=0.02). As such, while the observed genetic and antigenic properties of H3N2/K were not of themselves significantly outside of the expectation for year-on-year changes, the forecast antigenic change was unusually low given the genetic distinctiveness of the substrain. Selection for reasons other than antigenic novelty may have contributed to the global sweep of H3N2/K.

**Figure 3:**
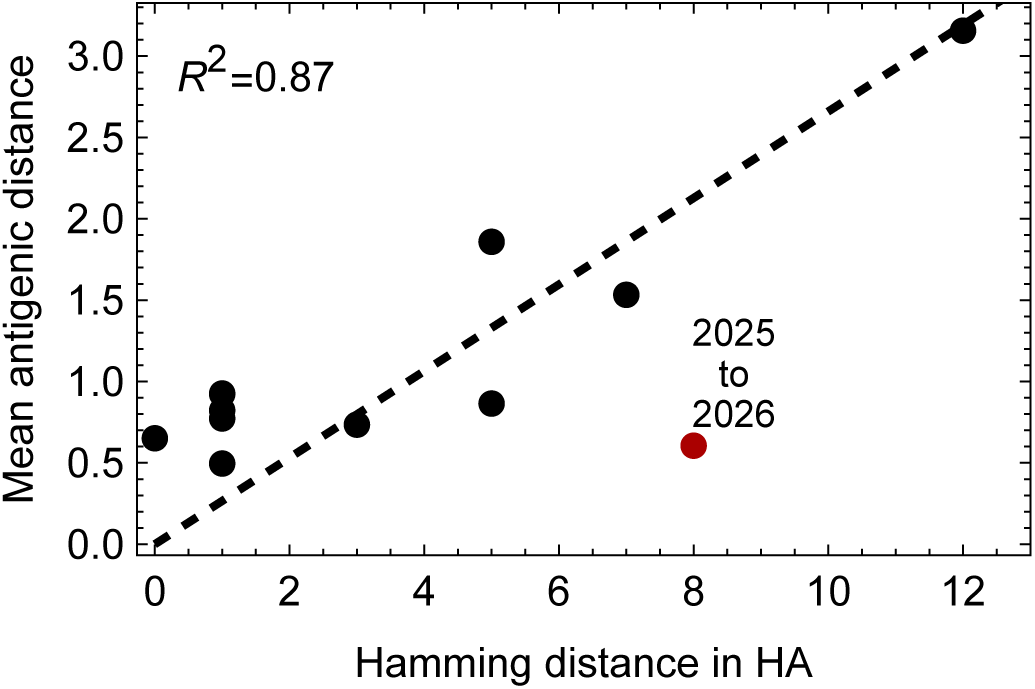
Relationship between the season-to-season genetic distance between consensus HA protein sequences and forecast antigenic distance under the PLANT model. Linear regression model fitted to genetic and antigenic distances calculated for year-to-year changes in the influenza A/H3N2 virus up to the 2024/25 season. The labelled red dot, showing the data for 2025/26, was not included in the model fitting. Relative to the model, this data point was a significant outlier.

### Antigenic properties of the H3N2/K strain: Serological analysis

Neutralisation assays performed on 486 serum samples derived from the NHS Greater Glasgow & Clyde adult patient population showed a significant decrease in neutralising ability for 2024-J and 2025-K relative to the 2022-Vacc strain (Figure 4). Samples were collected in May 2025, at the end of the 2024/5 influenza season, for which the 2022-Vacc strain formed part of the Northern Hemisphere vaccine recommendation^28^. Neutralisation levels differed in a manner dependent upon age, sex, and vaccination history. Once these were accounted for, an 18.7% decrease in overall neutralising activity for the 2024-J strain relative to 2022-Vacc, and a 20.4% decrease in the relative overall neutralising activity for 2025-K (Figure 4A). No overall differences in neutralisation were observed between 2024-J and 2025-K (p=0.66). However, antigenic differences between those two virus subclades appeared when a stratified analysis was performed. For example, the neutralisation of 2024-J was weaker than that of 2025-K among individuals with no record of vaccination (Figure 4D, p=0.032). More recent vaccination increased virus neutralisation, with a stronger effect on 2024-J strain. Thus, among individuals vaccinated in the 2024-25 season, the order was reversed, with 2024-J being more strongly neutralised than 2025-K (p=0.009): The latter was better able to escape immunity among the more recently vaccinated.

**Figure 4:**
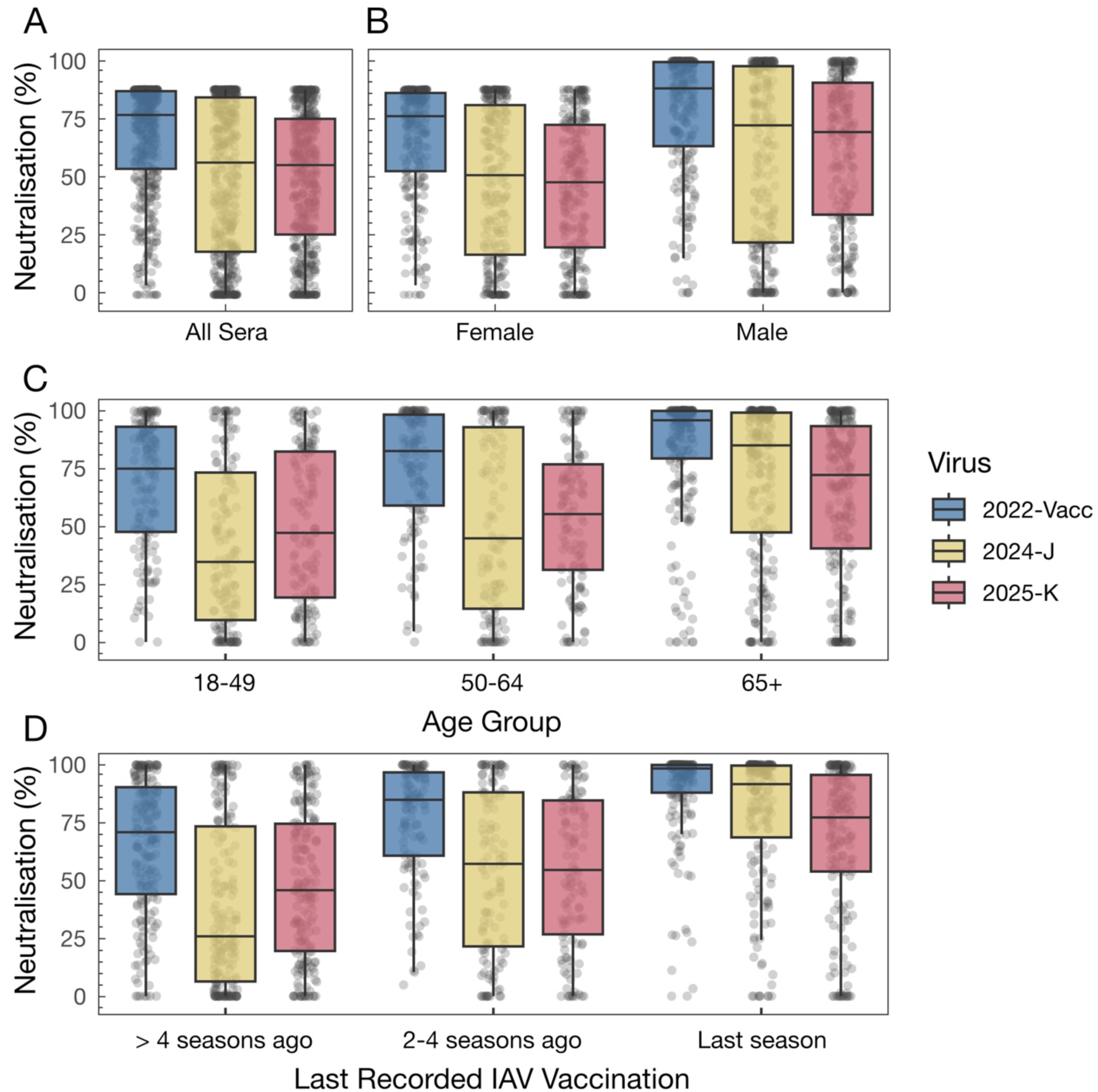
Neutralising activity of serum samples. Boxplots show percentage neutralisation of recombinant influenza viruses harbouring the surface glycoproteins of ENG2025 (H3N2/2025-K), ENG2024 (H3N2/2024-J), and the 2024/5 vaccine strain H3N2/2022-Vac. Data are stratified by **A.** Virus, measured across all indiviudals. **B.** Virus and sex; **C.** Virus and age group; **D.** Virus and vaccination history. These stratifications reflect all significant first-order effects and interactions identified through forward model building using beta regression.

Other patterns in neutralising activity were detected, with a small but significant increase of 3.1% in neutralisation for sera from males rather than females (Figure 4B, p=0.0261). People aged 65+ had a 5.5% increased neutralisation relative to people aged 18-49 (p=0.0152). The apparently larger differences between age groups observable in Figure 4 were explained in our analysis by differing levels of vaccination, with older people tending to be vaccinated more frequently. First-order differences between groups arising from our analysis are shown in Table 1, with further details provided in Supplementary Tables S3 to S5.

**Table 1:** Model-estimated first-order contrasts in serum neutralising activity. Values are predicted differences in percentage neutralisation (Reference − Comparison) from a beta regression model, averaged over all other covariates with equal weighting. P values are Tukey-adjusted for multiple comparisons.

| Variable | Reference | Comparison | Estimate | SE | p value |  |
| --- | --- | --- | --- | --- | --- | --- |
| **Virus** | 2022-Vacc | 2024-J | -18.7% | 1.7% | **<0.0001** | **\*\*\*** |
|  | 2022-Vacc | 2025-K | -20.4% | 1.7% | **<0.0001** | **\*\*\*** |
|  | 2024-J | 2025-K | -1.6% | 1.9% | 0.6614 |  |
| **Sex** | Female | Male | 3.1% | 1.4% | **0.0261** | **\*** |
| **Age Group** | 18-49 | 50-64 | 3.6% | 2.1% | 0.2139 |  |
|  | 18-49 | 65+ | 5.5% | 2.0% | **0.0152** | **\*** |
|  | 50-64 | 65+ | 2.0% | 1.8% | 0.5120 |  |
| **Time Since Last**<br>**IAV Vaccination** | > 4 seasons | 2-4 seasons | 7.4% | 1.9% | **0.0005** | **\*\*\*** |
|  | > 4 seasons | 1 season | 18.9% | 1.9% | **<0.0001** | **\*\*\*** |
|  | 2-4 seasons | 1 season | 11.5% | 2.1% | **<0.0001** | **\*\*\*** |

Neutralising activity was positively correlated between all three pairs of virus strains (2022-Vacc vs 2024-J R^2^ = 0.78; 2022-Vacc vs 2025-K R^2^ = 0.53; 2024-J vs 2025-K R^2^ = 0.47). The weaker correlations involving 2025-K suggest that the response to the K subclade involved antibody responses more distinct than those induced by the vaccine strain, without those responses necessarily being weaker in effect than those to subclade J.

### CD8+ T-cell based immune escape

While CD8+ T cell-driven immune escape has the potential to contribute to virus evolution, a computational analysis found no clear evidence relative to an HLA panel collected from the UK population^29^, favouring escape associated with the H3N2/K substrain (Figure 5). The forecast extent of CD8+ T Cell escape was calculated using the netMHCpan software package for each of a set of variants associated with the emergence of H3N2/K substrain, and for an opportunity space comprising all nonsynonymous mutations to the J subclade influenza virus A/Scotland/005_7/2024. Calculated scores were converted to Z-scores. The mean Z-score for an H3N2/K variant was slightly higher than that for the opportunity space, at 0.18 compared to - 0.02, in the direction of increased escape. However, comparison of the distributions of Z-scores suggested these were not significantly different (p=0.06, Kolmogorov-Sminnov test).

**Figure 5:**
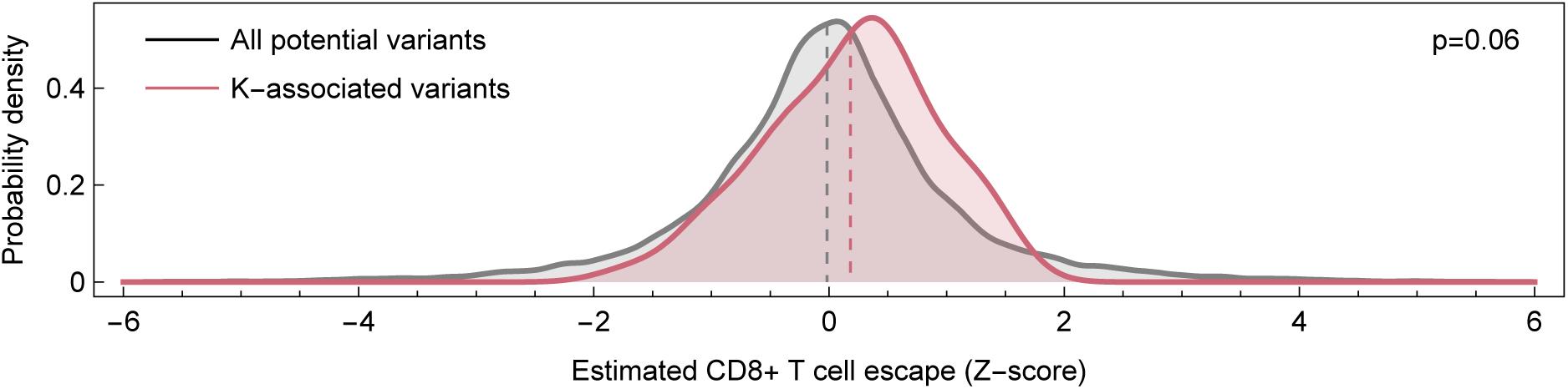
Estimated CD8+ T cell escape associated with the emergence of the H3N2/K subclade. Distributions were fitted to estimates of CD8+ T cell escape calculated relative to a distribution of HLA alleles representative of the UK population. Vertical dashed lines show the mean of each distribution.

## Discussion

Here we have combined different computational and experimental tools to evaluate the H3N2/K subclade in its historical and evolutionary context. Where computational approaches allowed for an easy comparison of historical strains, and a setting of results in context, serological measurements provided a specific and direct insight into the extent of immune escape provided by H3N2/K in its contemporary context. Both approaches agreed on the low antigenic difference between H3N2/K and its immediate predecessor. Our results from the PLANT computational model present the subclade as something of an outlier, breaking a historically strong pattern of correlated genetic and antigenic change.

Our results highlight differences in assessments of antigenic distinctiveness arising from serological analysis of samples from humans and ferrets, suggesting the value of human sera for obtaining an accurate picture of human immune responses. While the ferret model is well established for studying influenza infection^30^, and has long been used to assess antigenic changes between virus strains^31,32^, the immune response of a previously naïve animal does not necessarily capture the breadth of immune response accumulated over a lifetime in humans^33–35^: This lack of an immune history is a known limitation for assessing viral antigenic properties^36^. In this context, rapid assessment of viral samples using contemporary human sera could provide a more direct insight into the likely impact of an outbreak. Where early assessments of viral strains contribute to public health messaging, the importance of accuracy in assessments is particularly heightened.

One feature of note in our analysis is the breakdown of responses according to age and vaccination status. Clinical outcomes of influenza infection are stratified by age, with infection having generally more severe consequences in elderly people^37^. Where individuals in a population have distinct serological responses to a strain, the response of more vulnerable individuals may be the key determinant of the public health impact of a strain. Measurements of serological response that are stratified by age and vaccine status may therefore provide an improved picture for forecasting than simple population-average responses.

Our results highlight a distinction between measures of antigenic distinctiveness obtained in a small number of specific experiments carried out using samples from ferrets, and those obtained via machine learning, trained non-specifically upon a large set of ferret data. In this case, in describing the antigenic distinctiveness of the J and K strains, the PLANT model produced an outcome qualitatively closer to the human serological data than to the direct experimental measurement of antigenic escape in ferrets. The PLANT model had no direct knowledge of the K subclade, being trained on data only up to the year January 2024. A potential interpretation of this result is that the machine learning approach, in aggregating a large amount of data, evens out noise that can arise in small-scale animal experiments. Parallel assessments of ferret data, combining individual measurements of strain-specific immunity in animals with calculations generated from the context of machine-learning approaches, could enhance our understanding of immune response, the machine-learning model indicating the consistency, or lack of consistency, of an experimental result with our prior knowledge about patterns of viral immune escape.

As highlighted in this work, forecasting the impact of influenza epidemics remains an important and challenging task. While the drivers of influenza seasonality are well known, encompassing both social and environmental factors^38,39^, further work may be required on the specific attribution of an early or late season to changes in viral infectivity: Timing alone is insufficient as a metric of outbreak severity. While ferret data has for a long time represented the state of the art in quantifying human immune responses to influenza, novel approaches that incorporate human serological data might improve interpretation, potentially provide more accurate alternatives to epidemiological forecasting. Further work assessing the potential of these approaches to refine scientific understanding, and contribute to objective public health messaging, is a priority for future research.

## Methods

### Influenza surveillance data

Weekly laboratory-confirmed influenza virus infections from individuals presenting to General Practitioners with respiratory symptoms between October 3^rd^, 2022 and January 12^th^, 2026, were obtained from the Public Health Scotland (PHS) Community Acute Respiratory Infection (CARI) surveillance dataset^40^. Positive test rates for influenza A infection were calculated from these data.

### Sequence dataset assembly and phylogenetic analysis

Influenza A virus sequences were retrieved from the NCBI Entrez databases using the taxonomic identifier and an in-house Python tool interfacing with the E-utilities API, and supplemented with sequences from GISAID (all records available as of 12 February 2026). Sequences were restricted to H3 subtype HA coding sequences from human hosts. Records failing quality control were excluded: HA length < 1680 nucleotides (excluding N and non-IUPAC characters), atypical genetic divergence, or inconsistent host annotations. Redundant records were identified by strain name and removed to minimize duplication across databases. To reduce dataset size while preserving diversity, sequences were clustered with MMseqs2 (v15.6f452)^41^ at a 0.988 sequence identity threshold. Cluster representatives sequences were aligned using MAFFT (v7.453)^42^ with default parameters and a maximum-likelihood phylogeny was inferred with IQ-TREE (v2.4.0)^43^ under the best-fit substitution model. The clade 3C and descendant lineages were identified from this global tree and the relevant clusters expanded to include a non-redundant set of clade 3C sequences. IQ-TREE was used to reconstruct a comprehensive tree for the clade 3C and descendant linages and finally treetime (v0.10.0)^44^ was used to reconstruct the ancestral amino acid changes along the backbone of the phylogeny.

### Sequence analysis

In order to assess the timings of changes in H3N2 evolution, aligned protein sequences were split by the year in which they were collected, identifying the consensus sequence from each year. Mutations identified in the backbone of the phylogenetic tree were assigned to times according to the year in which the corresponding change in the consensus sequence was observed. This metric reflects the time at which mutations became dominant in the viral population. A semi-quantitative metric was used to assess numbers of mutations occurring each year, using a proxy measure based upon statistical significance. The mean number of nonsynonymous substitutions per year was calculated, using this to characterise a Poisson distribution of the expected number of mutations per year. Factors such as genetic linkage mean that the process of variant fixation in influenza populations is highly complex, and vulnerable to peaks^45^. In this context, our approach provides a benchmark indication of higher versus lower numbers of fixations per year, short of a formal measure of statistical significance.

### Antigenic escape modelling

The sequence lineage was embedded using PLANT^27^ to obtain 3D antigenic coordinates describing its relative position to other H3N2 sequences. The PLANT model trims sequences to consider just the HA1 region of haemagglutinin, and therefore neglects the impact of changes within HA2 or neuraminidase. The mean position of strains from each year was calculated within this antigenic projection, before calculating Euclidean distances between mean annual positions and specific viral strains.

### Analysis of outliers in genetic and antigenic data

Data from seasons prior to the emergence of H3N2/K was used to parameterise distributions of the number of nonsynonymous substitutions in the HA protein per year, and of the mean forecast antigenic distance between viruses each year according to the PLANT model. Changes in the 2025/26 season were then compared to these distributions.

For the combined analysis of genetic and forecast antigenic data, a linear regression model, restricted to pass through the origin, was fitted to data from prior to the 2025/26 season, representing the antigenic distance as a function of the genetic distance. We then used an external studentised residual test to evaluate the probability that the 2025/26 value was an outlier with respect to this model^46^.

### CD8+ T cell escape modelling

The CD8scape package^47^, implementing netMHCpan^48^, was used to assess the potential extent of CD8+ T cell escape in two sets of viral mutations. A set of all potential nonsynonymous mutations was generated across the eight viral segments of the virus A/Scotland/005_7/2024. A second list of mutations associated with the K subclade was also identified:

CD8+ T cell escape occurs in the context of specific HLA haplotypes. Here the distribution of haplotypes from the Anthony Nolan register^29^ was used to provide an approximate picture of escape acting on the level of the UK population, assigning frequencies to haplotypes observed in the register. For each haplotype *h*, a weight *w_h_* was assigned, corresponding to its frequency within the register.

Each change in the HA backbone was then evaluated using CD8scape relative to this panel. For each amino variant, CD8scape generated sets of all ancestral and derived peptides of length between 8 and 11 amino acids, spanning the variant. For each the ancestral and derived sets, we calculated values r_a,h_ and r_d,h_ respectively, describing the highest rank obtained by a peptide in each set, calculated from netMHCpan4.2, when considered against a reference set of peptides, versus the haplotype *h*. We now calculated the harmonic mean statistics

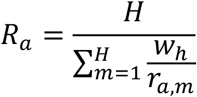

and

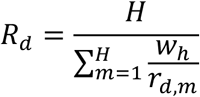

Where H is the number of haplotypes. The escape parameter was then calculated as a log fold change

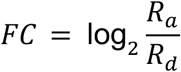

### Serum samples

Serum samples were provided by the National Health Service Greater Glasgow and Clyde (NHSGGC) Biorepository. Random residual biochemistry serum samples (n = 486) from primary (general practices) and secondary (hospitals) health care settings were collected by the NHSGGC Biorepository. Associated metadata, including age, sex, date of sample collection, and date of influenza vaccination were provided by the NHSGGC Biorepository.

### Viruses

Viruses were rescued by reverse genetics as previously described^49^. Viruses carried the internal genomic segments of A/Puerto Rico/8/34 (PR8) and the surface glycoproteins (HA and NA) of A/England/01837755/2025 (H3N2/2025-K, GISAID accession EPI_ISL_20210731); A/England/5160964/2024 (H3N2/2024-J, GISAID accession EPI_ISL_19667581) or A/Thailand/8/2022 (H3N2/2022-Vac, GISAID accession EPI_ISL_14991375) Plasmids encoding the internal genomic segments of PR8 were a kind gift of Prof. Ron Fouchier^50^. Plasmids encoding the HA and NA of H3N2/2022-Vac, H3N2/2024-J and H3N2/2025-K were synthesised and cloned by GeneArt Thermo Fisher.

### Serology assays

MDCK-SIAT cells were seeded at a density of 3 x10^5^ cells/ml ∼24h before infection. Serum samples were diluted 1: 75 in DMEM (Thermo Fisher Scientific Life Technologies 31966021) with 0.25% bovine serum albumin (BSA) (Thermo Fisher Scientific Life Technologies 15260037). 1.2 x10^3^ plaque forming units (PFU) of either: H3N2/2025-K, H3N2/2024-J, H3N2/2022-Vac was added 1:1 to each serum dilution in triplicate. The mixtures were incubated at 37 °C for 1 h and 100 µl was added to the MDCK-SIAT cells. The untreated virus input was 1.2 x10^3^ plaque forming units (PFU) mixed 1:1 with DMEM 0.25% BSA. Polyclonal anti-sera raised in sheep against the HA of A/Cambodia/e0826360/2020(National Institute of Biological Standards and Controls: 21/118) was used as a positive neutralization control. Plates were incubated at 37 °C, 5 % CO_2_ for 16 h, media removed and fixed in 10% formalin. Cells were permeabilized with 1 % Triton-X for 10 min. Virus positive cells were detected using a monoclonal antibody raised against IAV nucleoprotein (NP) (European Veterinary Labs, Clone EBS-I-238) at a dilution of 1: 1500 and then a secondary antibody (goat anti-mouse IgG Alexa Flour 488, A-11001) also at 1: 1500. Positive cells were counted using a Nexcelom Celigo imaging cytometer. For each plate, the mean total count for the input control was calculated. The total virus- positive cell count for each sample was then expressed as a percentage of the mean input count. These values were subtracted from 100, to give a percentage neutralization value. Values with negative neutralization values, i.e. higher counts than the input, were set to zero.

### Analysis of serology data

Neutralisation data were analysed as proportions using beta regression, adjusted using a Smithson-Verkuilen transformation to avoid exact values of 0 or 1^51^. A forward model building approach was used, beginning with a null (intercept only) structure and assuming constant precision. Plausible explanatory variables – virus (H3N2/2025-K, H3N2/2024-J, H3N2/2022-Vac); vaccination recency (last vaccinated in the 2024-2025 influenza season, last vaccinated in the 2021-2024 influenza seasons, no record of vaccination across the 2021-2025 influenza seasons); age group (18-49, 50-64, 65+); sex; and experimental block – were evaluated sequentially. Model extensions were retained if Akaike information criterion (AIC) indicated sufficient support for the increased model complexity. Additions to the precision model were considered before additions to the fixed-effects model, and first order effects were evaluated before second- and then third-order interaction terms. This resulted in a precision model with first-order effects of virus and experimental block, and a fixed-effects model with all possible first-order effects and two second-order interactions: one between virus and vaccination recency, and another between vaccine recency and age group. Effects of individual explanatory variables are reported as estimated marginal means on the response scale, averaged over the levels of other covariates with equal weighting, and the statistical significance of pairwise comparisons was assessed using asymptotic (Wald) z-tests. Correlations in neutralisation were assessed using Pearson correlation coefficients calculated on logit-transformed neutralisation values. All analyses were performed in R version 4.5.2 using the betareg (v3.2.4,^52^) and emmeans (v1.11.1, ^53^) packages.

## Ethics

Ethical approval for the collection of residual samples from Clinical Biochemistry was provided by NHSGGC Biorepository (application 837).

## Code and Data availability

All data and analysis scripts used in this study can be found on GitHub: https://github.com/ryanmimrie/Publications-2026-H3N2-K-Cross-Neutralisation.

## Acknowledgements

Funding to EH, DLR and PRM from the Medical Research Council (MRC) and Department for Environment, Food and Rural Affairs (Defra, UK) as FluTrailMap-One Health [MR/Y03368X/1]; funding from the Medical Research Council (MRC) to the MRC-University of Glasgow Centre for Virus Research to EH, BJW, ET, DLR, CJRI and PRM as MC_UU_0034/1, MC_UU_0034/2, MC_UU0034/3, MC_UU_00034/5 and MC_UU_00034/6. A record of viral genome sequences used in this study, acquired through GISAID, is provided here: https://doi.org/10.55876/gis8.260501cz

## Author Contributions

Conceptualization: KD, RMI, ET, CJRI, PRM; Data curation: KD, RMI, OAM, LM, EWS, ZW, JH; Formal analysis: KD, RMI, OAM, KL, DLR, CJRI, PRM; Funding acquisition: EH, BJW, ET, DLR, PRM; Investigation: KD, RMI, OAM, LM, EWS, SR, KL, HC, VS, ZW, SKW, JZ, JH, CJRI; Methodology: KD, RMI, EWS, JH, DLR, CJRI, PRM; Project administration: DLR, CJRI, PRM; Resources: DLR, CJRI, PRM; Software: RMI, OAM, EWS, KL; Supervision: EH, BJW, JH, DLR, CJRI, PRM; Validation: KD, RMI, JH, CJRI; Visualization: RMI, OAM, CJRI; Writing – original draft: KD, RMI, CJRI, PRM; Writing – review & editing: KD, RMI, EH, JH, DLR, CJRI, PRM

