## Supplementary Information for "Understanding the emergence of the influenza A/H3N2 K subclade in its historical and evolutionary context"

**Supplementary Figures**

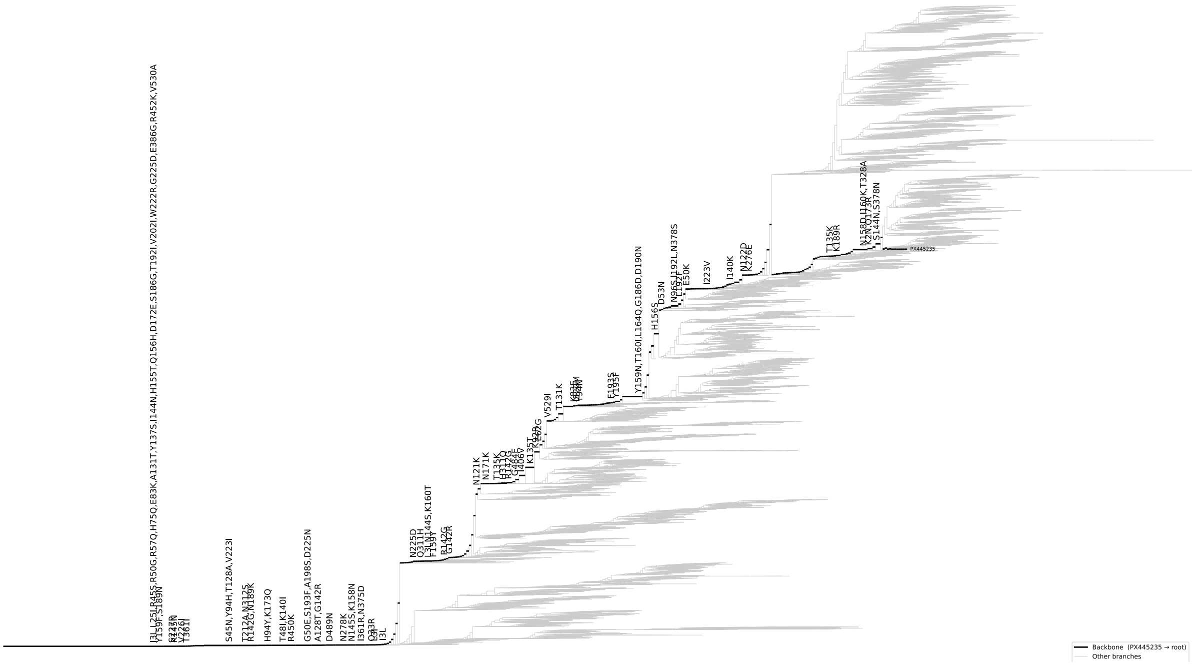

**S1 Figure: Phylogenetic tree detailing substitutions in the influenza A/H3N2 lineage.** The tree comprises information from 11480 sequences, representing the recent history of the HA segment of influenza A/H3N2. Non-synonymous substitutions in the tree leading to the H3N2/K substrain are shown on the backbone of the tree.

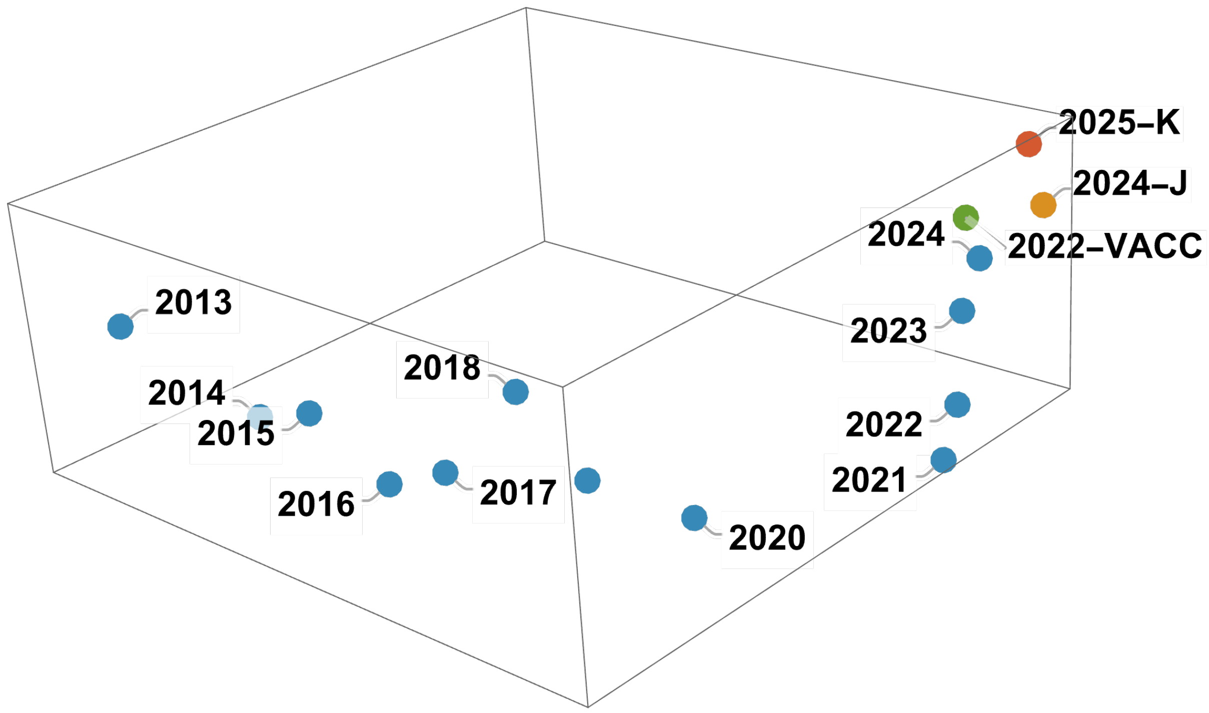


**S2 Figure: Mapping of HA1 sequences into predicted antigenic space achieved using the PLANT model.** Distances between dots show the extent of the predicted antigenic difference between strains. Blue dots represent the average positions of sequences from historical influenza seasons. The green point 2022-VACC indicates the HA1 sequence of the 2022 vaccine strain. The orange point 2024-J describes the location of a representative sequence from the 2024 H3N2 J subclade. The red point 2025-K describes the location of a representative sequence from the 2025 H3N2 K subclade.

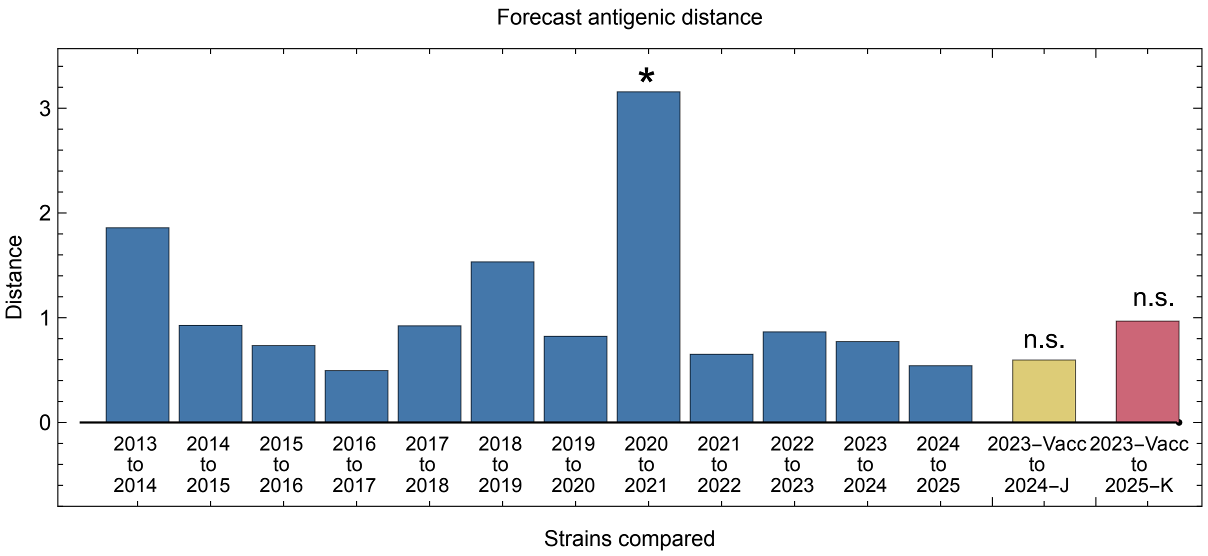


**S3 Figure: Forecast antigenic distances including the 2025/6 vaccine strain**.

The 2025/6 winter vaccine strain A/Croatia/10136RV/2023 is here denoted 2023-Vacc. Forecast antigenic distances between this and the representative sequences from clades J and K are shown in yellow and red. Other bars show mean forecast antigenic distances between sequences between sequences from previous seasons.

### Supplementary Tables

**S1 Table: Distributions fitted to numbers of year-to-year non-synonymous substitutions in the HA protein.** A negative binomial distribution (highlighted in italics) provided the best fit to the data.

| **Distribution** | **Likelihood** | **Bayesian Information Criterion (BIC)** |
|---|---|---|
| Poisson | -34.0773 | 70.6395 |
| *Negative binomial* | *-27.4059* | *59.7816* |

**S2 Table: Distributions fitted to mean year-to-year changes in forecast antigenic position.** A lognormal distribution (highlighted in italics) provided the best fit to the data.

| **Distribution** | **Likelihood** | **Bayesian Information Criterion (BIC)** |
|---|---|---|
| Weibull | -10.1111 | 25.0179 |
| *Lognormal* | *-8.07897* | *20.9537* |
| Gamma | -9.12215 | 23.0401 |

**S3 Table: Model-estimated second-order contrasts in serum neutralising activity for the interaction of virus strain and vaccination recency.** Values are predicted differences in percentage neutralisation (Reference – Comparison) from a beta regression model, averaged over all other covariates with equal weighting. P values are Tukey-adjusted for multiple comparisons.

| Variable | Stratum | Reference | Comparison | Estimate | SE | p value | |
|---|---|---|---|---|---|---|---|
| **Vaccination Group *within* Virus** | 2022-Vacc | > 4 seasons | 2-4 seasons | 7.9% | 2.7% | **0.0090** | ** |
| | | > 4 seasons | 1 season | 16.9% | 2.3% | **<0.0001** | *** |
| | | 2-4 seasons | 1 season | 9.0% | 2.4% | **0.0005** | *** |
| | 2024-J | > 4 seasons | 2-4 seasons | 11.1% | 3.5% | **0.0050** | ** |
| | | > 4 seasons | 1 season | 27.0% | 3.2% | **<0.0001** | *** |
| | | 2-4 seasons | 1 season | 15.9% | 3.5% | **<0.0001** | *** |
| | 2025-K | > 4 seasons | 2-4 seasons | 3.1% | 3.5% | 0.6501 | |
| | | > 4 seasons | 1 season | 12.7% | 3.3% | **0.0003** | *** |
| | | 2-4 seasons | 1 season | 9.6% | 3.6% | **0.0205** | * |
| **Virus *within* Vaccination Group** | > 4 seasons | 2022-Vacc | 2024-J | -23.1% | 2.8% | **<0.0001** | *** |
| | | 2022-Vacc | 2025-K | -17.3% | 2.8% | **<0.0001** | *** |
| | | 2024-J | 2025-K | 5.8% | 3.1% | 0.1363 | |
| | 2-4 seasons | 2022-Vacc | 2024-J | -20.0% | 3.4% | **<0.0001** | *** |
| | | 2022-Vacc | 2025-K | -22.2% | 3.3% | **<0.0001** | *** |
| | | 2024-J | 2025-K | -2.2% | 3.8% | 0.8280 | |

| | | | | | | | |
|---|---|---|---|---|---|---|---|
| | 1 season | 2022-Vacc | 2024-J | -13.1% | 2.4% | **<0.0001** | *** |
| | | 2022-Vacc | 2025-K | -21.5% | 2.5% | **<0.0001** | *** |
| | | 2024-J | 2025-K | -8.5% | 2.9% | **0.0090** | ** |

**S4 Table: Model-estimated second-order contrasts in serum neutralising activity for the interaction of age group and vaccination recency.** Values are predicted differences in percentage neutralisation (Reference – Comparison) from a beta regression model, averaged over all other covariates with equal weighting. P values are Tukey-adjusted for multiple comparisons.

| **Variable** | **Stratum** | **Reference** | **Comparison** | **Estimate** | **SE** | **p value** | |
|---|---|---|---|---|---|---|---|
| **Vaccination Group *within* Age Group** | 18-49 | > 4 seasons | 2-4 seasons | 3.2% | 3.5% | 0.6244 | |
| | | > 4 seasons | 1 season | 19.8% | 3.6% | **<0.0001** | *** |
| | | 2-4 seasons | 1 season | 16.5% | 4.5% | **0.0006** | *** |
| | 50-64 | > 4 seasons | 2-4 seasons | 13.5% | 3.3% | **0.0001** | *** |
| | | > 4 seasons | 1 season | 16.4% | 3.6% | **<0.0001** | *** |
| | | 2-4 seasons | 1 season | 2.9% | 3.3% | 0.6403 | |
| | 65+ | > 4 seasons | 2-4 seasons | 5.4% | 3.3% | 0.2396 | |
| | | > 4 seasons | 1 season | 20.4% | 2.5% | **<0.0001** | *** |
| | | 2-4 seasons | 1 season | 15.0% | 2.7% | **<0.0001** | *** |
| **Age Group *within*** | > 4 seasons | 18-49 | 50-64 | 1.3% | 3.0% | 0.9074 | |
| | | 18-49 | 65+ | 4.6% | 2.8% | 0.2286 | |

| Vaccination Group | | | | | | | |
|---|---|---|---|---|---|---|---|
| | | 50-64 | 65+ | 3.3% | 3.4% | 0.5875 | |
| | 2-4 seasons | 18-49 | 50-64 | 11.5% | 3.7% | **0.0054** | ** |
| | | 18-49 | 65+ | 6.7% | 3.9% | 0.2010 | |
| | | 50-64 | 65+ | 4.8% | 3.1% | 0.2823 | |
| | 1 season | 18-49 | 50-64 | 2.1% | 4.1% | 0.8635 | |
| | | 18-49 | 65+ | 5.2% | 3.4% | 0.2741 | |
| | | 50-64 | 65+ | 7.3% | 2.7% | **0.0204** | * |

**S5 Table: Model-estimated serum neutralising activity across demographic and vaccination strata.** Values are predicted percentage neutralisation of recombinant influenza viruses (mean ± SE) from a beta regression model, shown for each combination of virus, sex, age group, and time since last influenza vaccination.

| Virus | Sex | Time Since Last IAV Vaccination | Age: 18-49 | Age: 50-64 | Age: 65+ |
|---|---|---|---|---|---|
| **2025-K** | Female | > 4 seasons | 44.8% (±2.5%) | 46.1% (±3.3%) | 49.6% (±3.1%) |
| | | 2-4 seasons | 43.4% (±3.9%) | 55.8% (±3.2%) | 50.5% (±3.5%) |
| | | 1 season | 58.3% (±4.2%) | 55.9% (±3.6%) | 64.8% (±2.3%) |
| | Male | > 4 seasons | 48.2% (±2.6%) | 49.6% (±3.2%) | 53.0% (±3.1%) |
| | | 2-4 seasons | 46.8% (±4.0%) | 59.2% (±3.1%) | 54.0% (±3.5%) |
| | | 1 season | 61.6% (±4.3%) | 59.1% (±3.5%) | 67.8% (±2.1%) |

| | | | | | |
|---|---|---|---|---|---|
| **2024-J** | Female | > 4 seasons | 39.1% (±2.4%) | 40.4% (±3.2%) | 43.8% (±3.1%) |
| | | 2-4 seasons | 45.6% (±4.0%) | 58.0% (±3.3%) | 52.8% (±3.5%) |
| | | 1 season | 67.1% (±3.9%) | 64.8% (±3.4%) | 72.9% (±2.0%) |
| | Male | > 4 seasons | 42.4% (±2.6%) | 43.7% (±3.2%) | 47.2% (±3.1%) |
| | | 2-4 seasons | 49.0% (±4.1%) | 61.3% (±3.2%) | 56.2% (±3.6%) |
| | | 1 season | 70.1% (±3.9%) | 67.9% (±3.2%) | 75.5% (±1.9%) |
| **2022-Vacc** | Female | > 4 seasons | 62.5% (±2.2%) | 63.7% (±3.0%) | 66.9% (±2.7%) |
| | | 2-4 seasons | 67.1% (±3.4%) | 77.2% (±2.3%) | 73.2% (±2.7%) |
| | | 1 season | 81.0% (±2.7%) | 79.4% (±2.4%) | 84.9% (±1.3%) |
| | Male | > 4 seasons | 65.6% (±2.3%) | 66.8% (±2.8%) | 69.8% (±2.5%) |
| | | 2-4 seasons | 70.1% (±3.4%) | 79.5% (±2.1%) | 75.8% (±2.6%) |
| | | 1 season | 83.1% (±2.6%) | 81.6% (±2.2%) | 86.6% (±1.2%) |
